# A Conserved Mechanism of Cardiac Hypertrophy Regression through FoxO1

**DOI:** 10.1101/2024.01.27.577585

**Authors:** Thomas G. Martin, Dakota R. Hunt, Stephen J. Langer, Yuxiao Tan, Christopher C. Ebmeier, Claudia Crocini, Eunhee Chung, Leslie A. Leinwand

**Affiliations:** Department of Molecular, Cellular, and Developmental Biology, University of Colorado Boulder, Boulder CO; Department of Biochemistry, University of Colorado Boulder, Boulder CO; BioFrontiers Institute, University of Colorado Boulder, Boulder CO; Department of Kinesiology, University of Texas at San Antonio, San Antonio, TX

**Author notes:** Corresponding Author Leslie A. Leinwand, Ph.D. BioFrontiers Institute Jennie Smoly Caruthers Biotechnology Building, D354 3415 Colorado Ave Boulder CO, 80303.

## Abstract

The heart is a highly plastic organ that responds to diverse stimuli to modify form and function. The molecular mechanisms of adaptive physiological cardiac hypertrophy are well-established; however, the regulation of hypertrophy regression is poorly understood. To identify molecular features of regression, we studied Burmese pythons which experience reversible cardiac hypertrophy following large, infrequent meals. Using multi-omics screens followed by targeted analyses, we found forkhead box protein O1 (FoxO1) transcription factor signaling, and downstream autophagy activity, were downregulated during hypertrophy, but re-activated with regression. To determine whether these events were mechanistically related to regression, we established an *in vitro* platform of cardiomyocyte hypertrophy and regression from treatment with fed python plasma. FoxO1 inhibition prevented regression in this system, while FoxO1 activation reversed fed python plasma-induced hypertrophy in an autophagy-dependent manner. We next examined whether FoxO1 was implicated in mammalian models of reversible hypertrophy from exercise and pregnancy and found that in both cases FoxO1 was activated during regression. In these models, as in pythons, activation of FoxO1 was associated with increased expression FoxO1 target genes involved in autophagy. Taken together, our findings suggest FoxO1-dependent autophagy is a conserved mechanism for regression of physiological cardiac hypertrophy across species.

**HIGHLIGHTS:**

- Post-prandial cardiac remodeling in Burmese pythons is associated with dynamic regulation of FoxO1 and autophagy.
- Regression of fed python plasma-induced hypertrophy in mammalian cardiomyocytes is FoxO1-dependent.
- Activation of FoxO1-dependent autophagy is a conserved feature of physiological hypertrophy regression in mammalian models.

## INTRODUCTION

The heart is a highly plastic organ that can modify structure and function to meet physiological demands. However, insight into the broad scope of molecular mechanisms that govern cardiac plasticity remains limited. There are several notable examples of robust, reversible cardiac remodeling occurring in vertebrates including in fish with thermal acclimation^1–4^, birds flying at high altitudes^5–7^, and pythons following consumption of large prey^8–12^. Such unconventional model organisms represent unique platforms in which to discover the molecular biology underlying cardiac plasticity. Physiological cardiac remodeling has also been observed in mammals with exercise or pregnancy, where the heart grows in response to increased circulatory demands^13–15^. Unlike hypertrophy arising from pathological conditions (e.g., hypertension) such as that which occurs in heart disease, physiological hypertrophy is associated with unchanged or improved cardiac function and is completely reversible with removal of the stimulus^13^. The cellular and molecular underpinnings of physiological cardiac growth are well established: circulating growth factors activate phosphoinositide-3-kinase (PI3K)/Akt/mammalian target of rapamycin (mTOR) signaling in cardiomyocytes, which stimulates protein synthesis^13^. However, the molecular mechanisms that regulate the regression of cardiac hypertrophy are largely unknown.

Burmese pythons (*Python bivittatus)* employ an ambush hunting predation strategy that is associated with extended months-long fasts. We and others have previously shown that the python heart undergoes substantial hypertrophy within three days of ingesting large prey^8–11^. The impetus for post-prandial cardiac growth is the elevated metabolic activity of the python’s organs during digestion, which necessitates increased oxygen delivery to distal tissues^11^. Notably, the cardiac hypertrophy observed is not a consequence of increased fluid content^8,10^, but rather results from elevated protein synthesis downstream of Akt/mTOR, leading to muscle growth^8,9,16^. Equally impressive to the magnitude of post-prandial cardiac hypertrophy in Burmese pythons is the rate with which this new muscle mass regresses. Indeed, previous studies have shown that post-prandial remodeling is nearly completely regressed just 10 days after a meal^9,12,16^. The magnitude of hypertrophy and rapidity of its regression make the python an ideal model organism in which to identify the molecular factors that regulate cardiac structural plasticity.

In the present study, we show activation of the transcription factor forkhead box protein O1 (FoxO1) contributes to regression of physiological hypertrophy in pythons. Multi-omics analyses revealed that hypertrophy in pythons was associated with reduced FoxO1 activity and suppression of downstream autophagy gene expression. FoxO1/autophagy were re-activated during regression.

Treating cultured mammalian cardiomyocytes with fed python plasma recapitulated physiological hypertrophy, the regression of which was dependent on FoxO1. Moreover, we found FoxO1-dependent cell size regression required activation of autophagy. To examine whether mammals might employ a similar mechanism during reversible cardiac hypertrophy, we studied hypertrophy regression from aerobic exercise training and pregnancy in mice. In both models, FoxO1 activity was potently activated during regression, which coincided with increased expression of FoxO1 gene targets involved in autophagy. Our findings suggest that FoxO1 activation is a key factor driving regression of physiological hypertrophy that is conserved across species.

## RESULTS

### Post-prandial ventricular remodeling in the Burmese python is associated with dynamic regulation of FoxO1 and autophagy

To identify factors associated with the regression of cardiac mass in Burmese pythons, we performed bulk RNA sequencing on mRNA from fasted, 1-day post-fed (DPF), 4DPF, and 6DPF python ventricles. All pythons received rodent meals equivalent to 25% of their body mass. Based on our previous work showing regression of mass occurs across an approximately 1 week period from 3-10DPF^9^, 6DPF represents a peak regression timepoint where ventricular mass is actively being reduced. We therefore hypothesized that factors differentially expressed at 6DPF compared with 1DPF would include those that mediate regression. K-means clustering was used to group differentially expressed genes into different expression patterns across the feeding time course and revealed three clusters: 1. Gene expression increased during hypertrophy and then decreased during regression, 2. Gene expression did not change during hypertrophy but significantly increased during regression, 3. Gene expression decreased during hypertrophy and then increased during regression (**Fig. 1A-C**). The two most downregulated genes during digestion, *FBXO32* and *BNIP3* clustered into the third group. When we analyzed the differentially expressed genes in Cluster 3, we found that the most significantly enriched pathways were regulation of macroautophagy and forkhead box protein (FoxO) signaling (**Fig. 1D-E**). FoxOs are known to regulate expression of the aforementioned genes and are master transcriptional regulators of macroautophagy, which facilitates protein and organelle degradation via the lysosome^17^. Further analysis of Cluster 3 according to transcription factor enrichment of differentially expressed genes identified FoxO1-mediated gene expression as the top transcriptional pathway affected (**Fig. 1F-G**). As expected, Cluster 1 predominantly included genes involved in protein synthesis and protein transport (**Supplemental Fig. 1**). Cluster 2, which was expected to include regression promoting factors, was defined by cell cycle and cellular respiration genes (**Supplemental Fig. 1**). These pathways are not expected to be involved in reversal of muscle mass given their lack of involvement in protein homeostasis or atrophic pathways.

**Figure 1.**
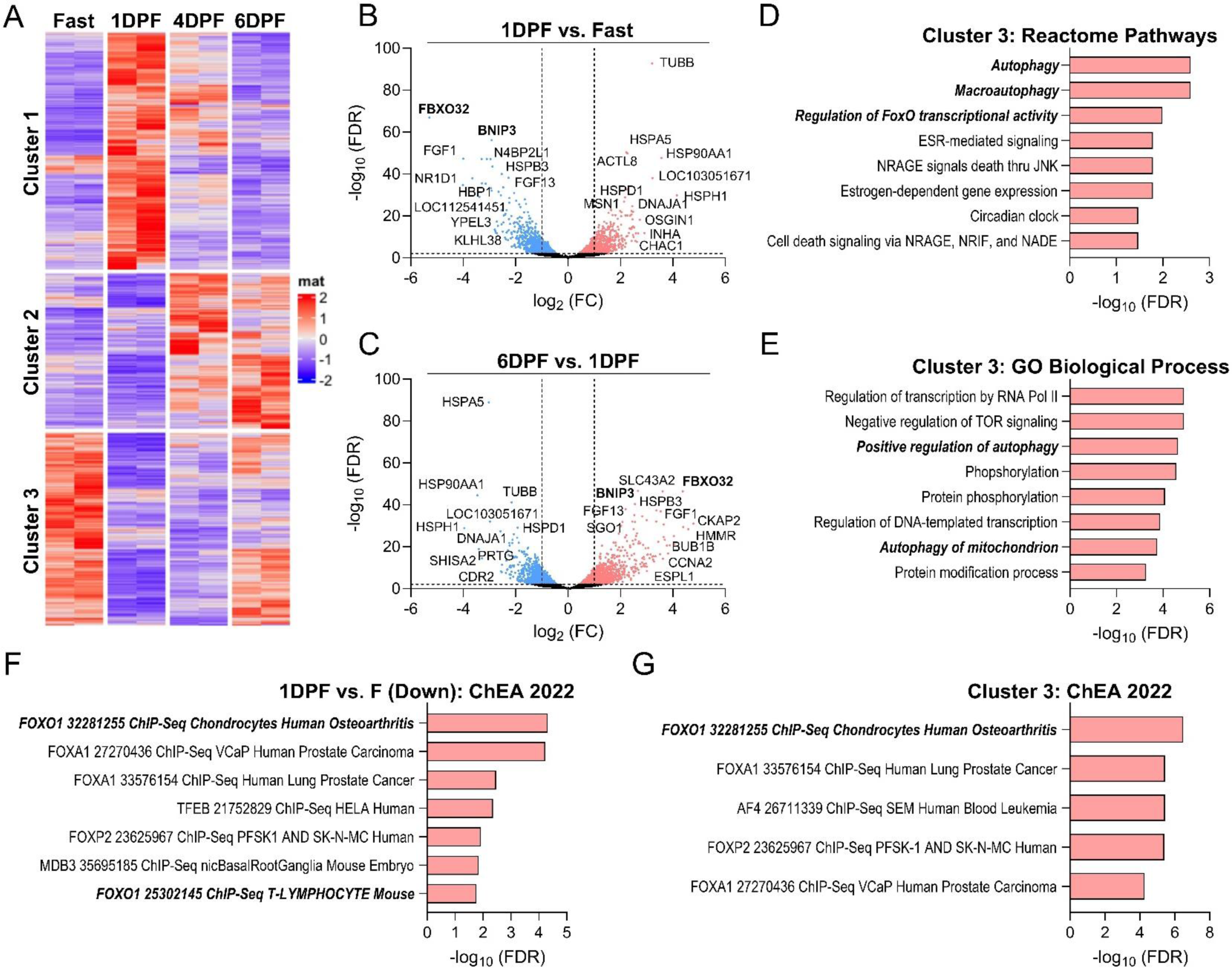
Post-prandial ventricular remodeling in the Burmese python is associated with dynamic regulation of FoxO1 signaling and autophagy. **A.** Heat map of differentially expressed python genes after feeding identified by RNA sequencing and partitioned into 3 clusters by K-means clustering. mat = matrix, representing the Z-score. **B.** Volcano plot 1DPF vs. Fasted gene expression highlighting FoxO gene targets *FBXO32* and *BNIP3* as the most significantly downregulated genes. **C.** Volcano plot of 6DPF vs. 1DPF indicating restored expression of FoxO target genes. D-E. Reactome pathway (D) and GO Biological Process (E) enrichment of differentially expressed genes in Cluster 3. F. Downregulated genes at 1DPF vs Fasted enriched by target genes of transcription factors in the ChEA 2022 database. G. Significantly differentially expressed genes identified in Cluster 3 in the RNAseq data enriched by target genes of transcription factors in the ChEA 2022 database. FDR = false discovery rate adjusted p-value, FC = fold-change.

To further characterize the molecular changes at the proteome-level, we performed relative quantitative proteomics analysis via tandem-mass-tag multiplexing of tryptic peptides from fasted, 1DPF, 4DPF, and 6DPF python ventricles. In total, 1,177 proteins were significantly differentially expressed (Adj. p ≤ 0.01, log_2_ fold-change ≥ 0.3) across the four groups (**Supplemental Fig. 2A**). The proteomics data further supported that dynamic changes in autophagy activity occur during digestion, as autophagy was reduced at 1DPF (increased expression of SQSTM1/P62) and then re-activated during regression at 6DPF (return of SQSTM1/P62 to fasted levels) (**Supplemental Fig. 2B-D**). Moreover, gene ontology analysis of the upregulated proteins at 6DPF identified proteins involved in macroautophagy and autophagosome maturation as significantly enriched compared to fasted (**Supplemental Fig. 2E**). Together, these multi-omics data indicate that FoxO1 and autophagy activity are dynamically modulated during ventricular remodeling in the python. Given the established role of these factors in regulating protein degradation in muscle^18,19^, and that muscle cell size and thus organ size is determined by the balance between protein synthesis and degradation, we hypothesized FoxO1/autophagy activation was the mechanism of regression in Burmese pythons.

### FoxO1 and autophagy are suppressed during hypertrophy development in Burmese pythons and re-activated with regression

To fully characterize the activity of FoxO1 and autophagy throughout digestion in pythons, we next conducted a larger Burmese python feeding study. Hearts were collected 0.5, 1, 2, 3, 6, and 10DPF and heart mass was normalized to pre-fed body weight. As previously observed^8,9,16^, we found that cardiac hypertrophy peaked around 2DPF and was completely regressed by 10DPF (**Fig. 2A**). This was preceded by activation of Akt at 1DPF, suggesting increased stimulation of protein synthesis (**Supplemental Fig. 3**). To examine FoxO1 activity, we measured phosphorylation of FoxO1 at threonine 24, which had previously been shown to cause cytosolic retention of FoxO1, rendering it transcriptionally inactive^20^. FoxO1 phosphorylation increased significantly by 0.5DPF and then started to decrease at 3DPF, reaching its lowest level at 6DPF (**Fig. 2B-C**). We next measured autophagy by LC3B expression and phosphorylation of the autophagy-initiating kinase ULK1. Serine 555 phosphorylation of ULK1 (indicating activation) and expression of the cleaved form of LC3B (LC3-II, marker of autophagosome maturation) were suppressed during hypertrophy and then increased with regression (**Fig. 2B, D-E**). Finally, we measured the expression of established FoxO1 target genes involved in autophagy: *BNIP3*, *CTSL*, *MAP1LC3B*, and *ULK1*. Transcript levels followed the same pattern as FoxO1 activity, decreasing significantly at 0.5-1DPF and then peaking at 6DPF (**Fig. 2F-I**). These data support that activation of FoxO1-dependent autophagy after peak digestion in pythons could underly the observed regression of ventricular mass back to baseline.

**Figure 2.**
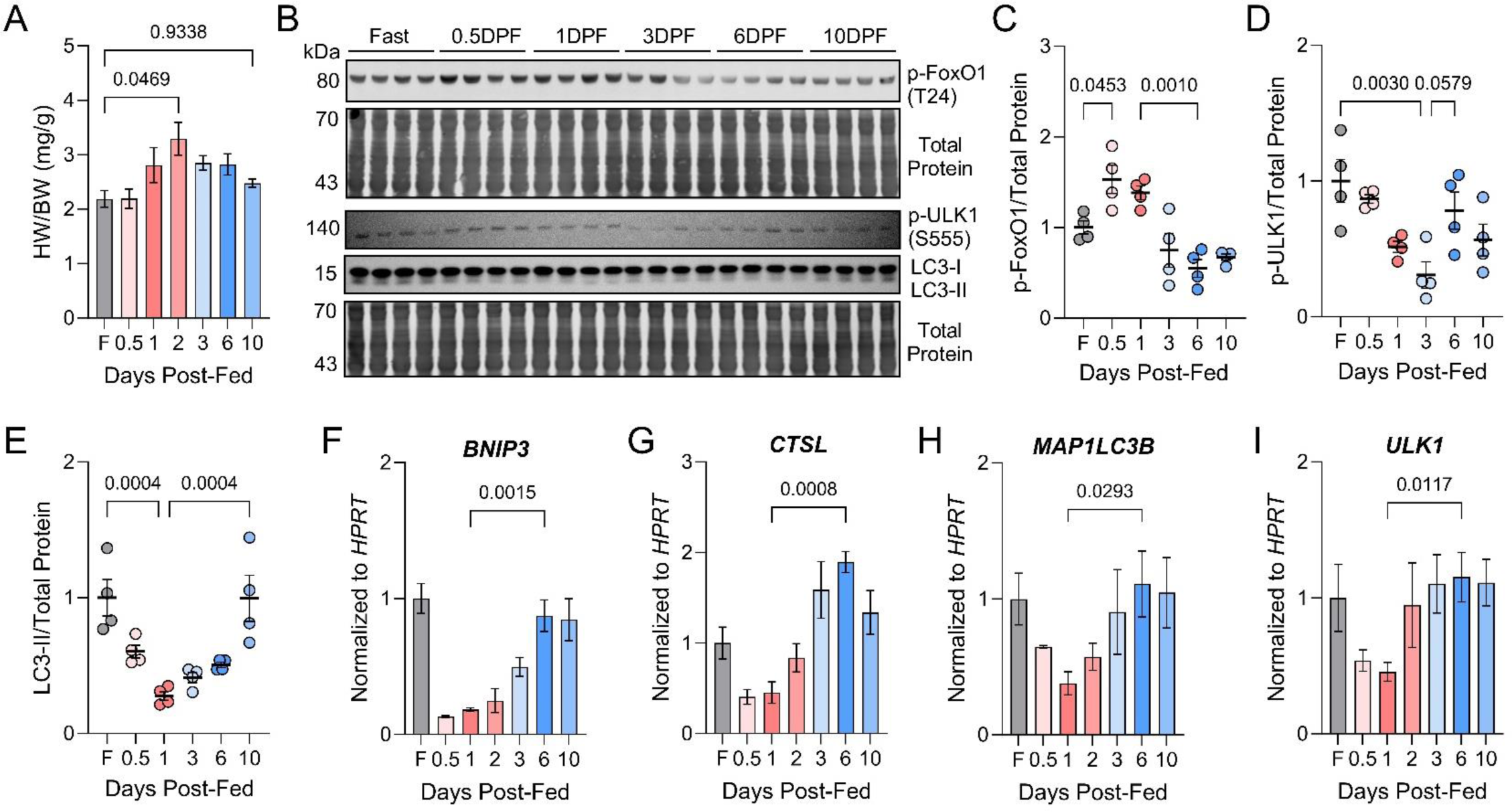
FoxO1 and autophagy activity are suppressed during hypertrophy development in the Burmese python and re-activated with regression. **A.** Python heart weight normalized to body weight throughout the feeding paradigm; F = fasted. **B.** Western blots for phosphorylated FoxO1, phosphorylated ULK1, and LC3B. **C-E.** p-FoxO1 (C), p-ULK1 (D), and LC3-II (E) expression normalized to total protein. **F-I.** qPCR analysis of the autophagy genes *BNIP3* (F)*, CTSL* (G)*, MAP1LC3B* (H)*, and ULK1* (I) normalized to *HPRT*. For all, n = 4 pythons/group and data were analyzed by one-way ANOVA.

### Physiological hypertrophy is recapitulated in NRVMs treated with fed python plasma

We had previously shown that cardiac hypertrophy in pythons was due to circulating growth factors in the plasma that increase during digestion^9^. Additionally, administration of fed python plasma (FPP) to neonatal rat ventricular myocytes (NRVMs) resulted in marked cellular hypertrophy^9^. In the current study, we adapted the NRVM system to mechanistically probe hypertrophy regression using a passive regression model. To model hypertrophy and regression, NRVMs were exposed to FPP for 24 hours (Treatment) and then the media was changed back to control media for 24-48 hours (Chase) (**Fig. 3A**). FPP, but not fasted python plasma, induced cellular hypertrophy matching the level initiated by the positive control phenylephrine (PE, 10µM), an α-adrenergic receptor agonist associated with pathological hypertrophy^21^. FPP-induced hypertrophy was rapidly reversible following plasma withdrawal (**Fig. 3B**). Importantly, FPP-treated cells displayed features of healthy remodeling, including eccentric hypertrophy and limited induction of fetal gene expression compared with PE (**Supplemental Fig. 4, Fig. 3C**). Protein synthesis rates, monitored by puromycin incorporation, increased with both PE and FPP, but returned to control levels during the chase. Fasted python plasma had no effect on protein synthesis (**Fig. 3D, Supplemental Fig. 4C**). As in the post-prandial python heart^9^, the increase in protein synthesis observed with FPP in NRVMs was preceded by activation of mTOR and Akt (**Fig. 3E-G**). Since protein synthesis rates were not significantly lower than control levels after removal of FPP, we hypothesized that activation of protein degradation must drive the regression of cell size. Therefore, we next measured autophagic flux and autophagy gene expression. We found that autophagy was activated six hours after withdrawing the hypertrophic agonists (**Fig. 3H-I**). Additionally, gene expression of *Bnip3, Map1lc3b,* and *Ulk1* increased following FPP removal (**Fig. 3J**), matching what was observed in the python ventricle during regression. These genes were not induced with removal of PE, suggesting there are distinct mechanisms of autophagy activation in the two settings.

**Figure 3.**
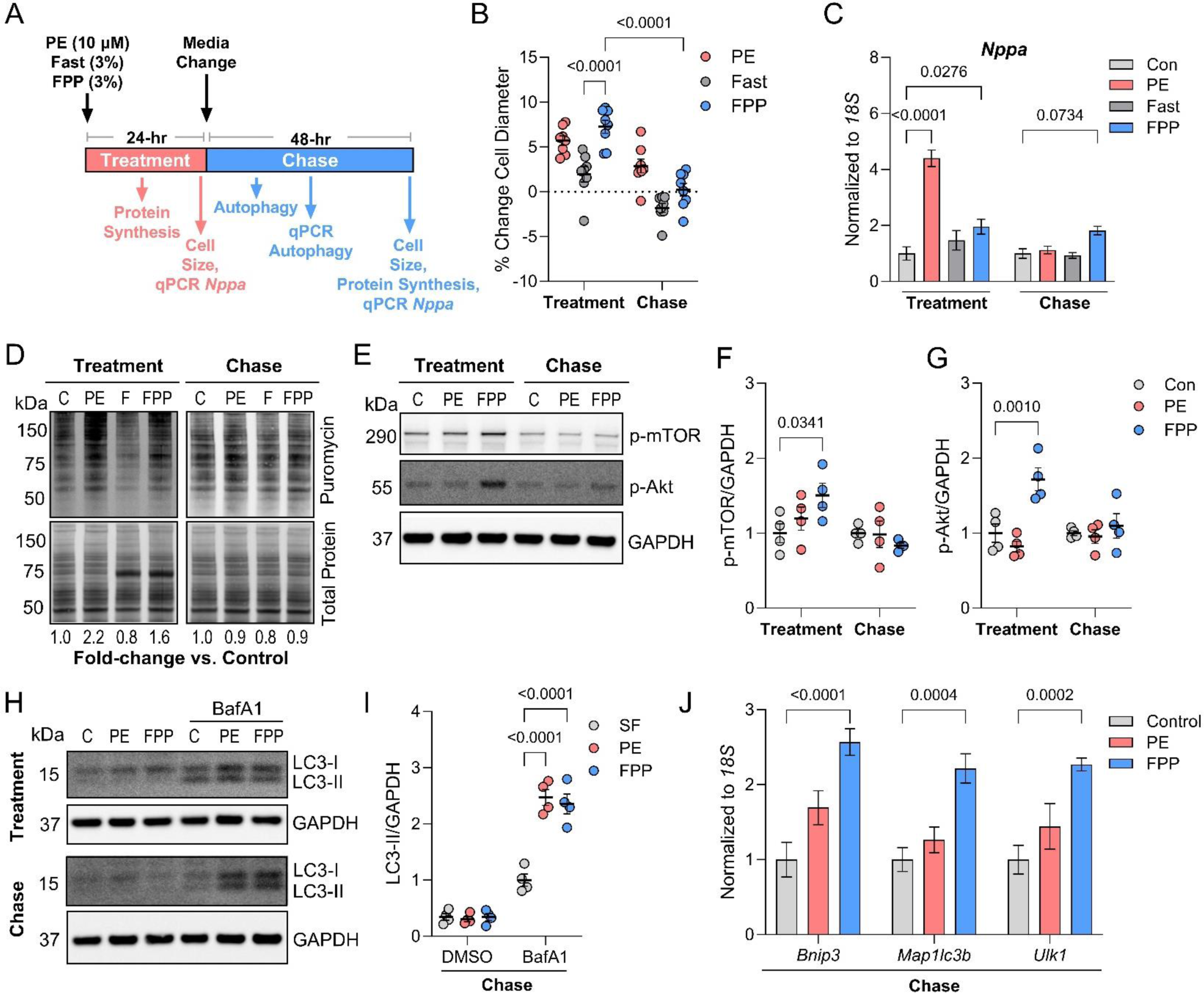
Mammalian cardiomyocytes treated with fed python plasma recapitulate molecular features of python post-prandial cardiac remodeling. **A.** Experimental paradigm. PE = phenylephrine, Fast = fasted python plasma, FPP = fed python plasma. **B.** Percent change in cell particle diameter compared with control for NRVMs treated with PE, 3% Fast, or 3% FPP during the Treatment and Chase. n = 10,000-30,000 cells/replicate, 8 biological replicates/group; two-way ANOVA. **C.** qPCR analysis for *Nppa* gene expression normalized to *18S;* n = 5/group, two-way ANOVA. **D.** Immunoblots for puromycin, representing protein synthesis rates. **E-G.** Representative immunoblots (E) and normalized expression of p-mTOR (F) and p-Akt (G); n = 4/group; two-way ANOVA. **H.** Representative immunoblot for LC3B in NRVMs during Treatment and Chase with and without BafA1. **I.** LC3-II expression normalized to GAPDH during the Chase. n = 4/group, two-way ANOVA. **J.** qPCR analysis for *Bnip3, Map1lc3b,* and *Ulk1* expression normalized to *18S* during the Chase. n = 5/group, two way-ANOVA. Con, C = control.

### FoxO1 inhibition prevents regression of fed-python plasma-induced hypertrophy

Given the association of hypertrophy regression in NRVMs with upregulation of autophagy gene expression and activity, we next measured FoxO1 activity by western blot for p-FoxO1. FPP treatment increased p-FoxO1 by ∼6-fold compared to control/PE and this inhibition of FoxO1 was rapidly reversible within 6 hours of agonist withdrawal (**Fig. 4A-B**). The effect of FPP on FoxO activity was unique to FoxO1, as FoxO3 phosphorylation was unaffected (**Fig. 4A**). Notably, regression in NRVMs after withdrawal of FPP was associated with nuclear translocation of FoxO1 (**Fig. 4C**), in agreement with increased transcriptional activity. To test whether FoxO1 was required for regression in NRVMs, a FoxO1-specific small molecule inhibitor (AS1842856) was added to the media during the chase. Cell size had not significantly regressed 24 hours after addition of AS1842856, while cells exposed to DMSO vehicle displayed near-complete regression (**Fig. 4D**). Examination of FoxO1-dependent gene expression at this time-point indicated that while the media change with vehicle resulted in induction of autophagy gene expression, AS1842856 completely abrogated this effect (**Fig. 4E-H**). Moreover, autophagic flux measured by LC3-II intensity in the presence of BafA1 was significantly reduced by AS1842856 (**Fig. 4I-L**).

**Figure 4.**
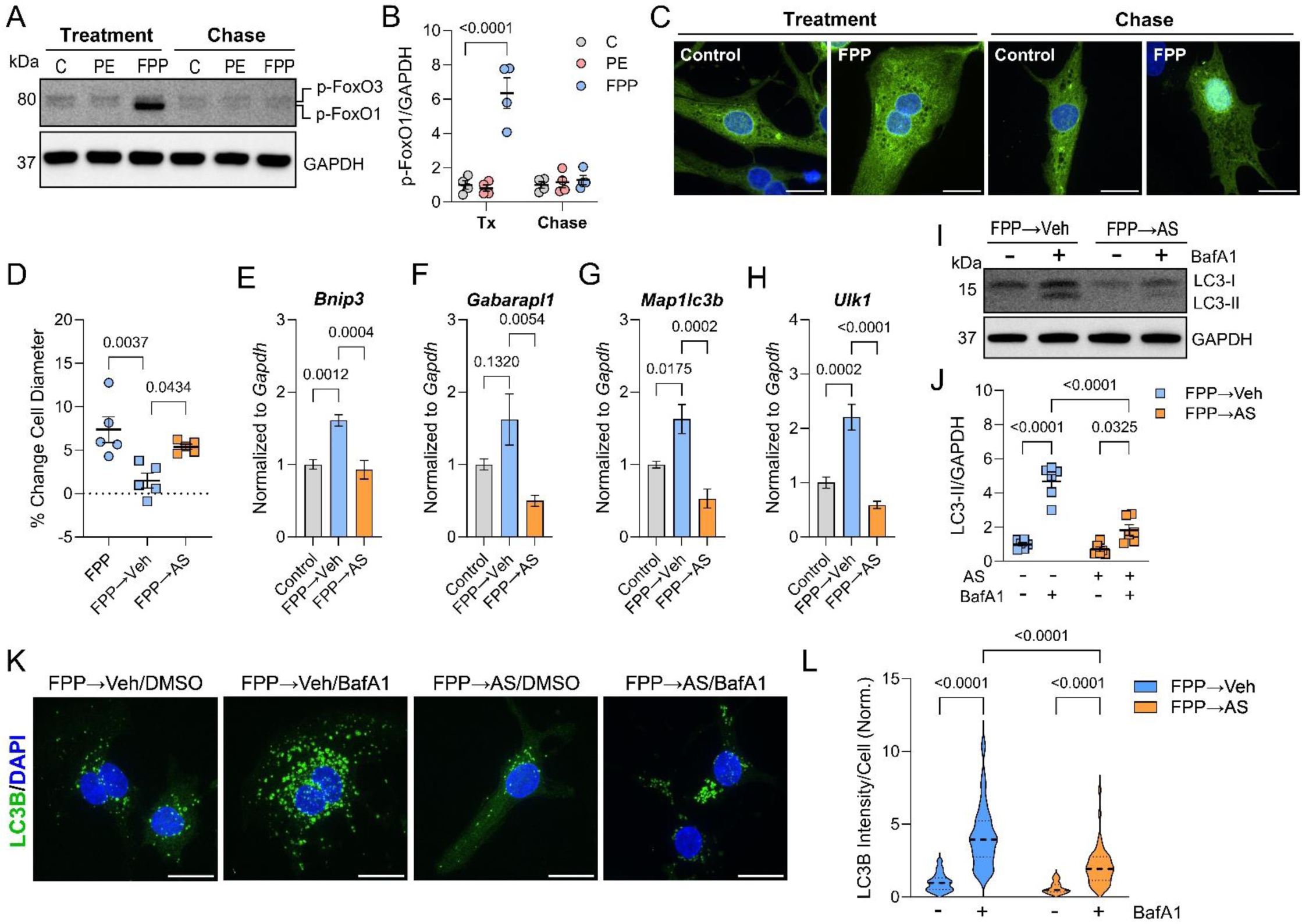
FoxO1 inhibition prevents regression of fed python plasma-induced hypertrophy. **A-B.** Representative immunoblot (A) and normalized expression of (B) phosphorylated FoxO1 (Threonine 24) during Treatment (24 hours) and Chase (6 hours post agonist withdrawal); n = 4/group; two-way ANOVA; C = control, Tx = treatment. **C.** Representative immunofluorescence microscopy images for NRVMs transduced with Ad-GFP-FOXO1_WT_ treated with FPP (Treatment) or 12 hours after withdrawal of FPP (Chase); 100X magnification, scale bar = 15 µm. **D.** Percent change in cell particle diameter compared with control for NRVMs treated with FPP, 24 hours after FPP removal with addition of DMSO vehicle or 2 µM AS1842856 (AS, FoxO1-specific inhibitor; n = 10,000-30,000 cells/replicate, 5 biological replicates/group; one-way ANOVA. **E-H.** qPCR analysis of autophagy genes *Bnip3* (E)*, Gabarapl1* (F)*, Map1lc3b* (G), and *Ulk1* (H) normalized to *Gapdh;* n = 6/group; one-way ANOVA. **I-J.** Representative immunoblot (I) and normalized expression of (J) LC3B 24 hours after FPP withdrawal and addition of AS; two-way ANOVA. **K-L.** Representative immunofluorescence microscopy images (K) and quantification (L) of LC3B-positive area (Green) in NRVMs transduced with Ad-GFP-LC3B after FPP withdrawal with or without the FoxO1 inhibitor AS1842856 and ± BafA1; 100X magnification, scale bar = 15 µm; n = 42 FPP→C/DMSO, 52 FPP→C/BafA1, 48 FPP→AS/DMSO, 49 FPP→AS/BafA1; two-way ANOVA.

### FoxO1 activation reverses fed python plasma-induced hypertrophy

Given the above result, we hypothesized that FoxO1 activation would promote regression in the presence of FPP. We therefore generated adenoviruses expressing constitutively active FoxO1 (Ad-FOXO1_CA_), FOXO1_CA_ in which the DNA-binding domain was deleted (Ad-FOXO1_ΔDB_), and an empty vector control. To test whether activation of FoxO1 could reverse FPP-induced hypertrophy, NRVMs were treated with FPP for 24 hours and then adenoviruses added into the FPP-containing media at 20 multiplicity of infection (MOI) and left for 24-48 hours (**Fig. 5A**). The two FoxO1 constructs displayed different localizations, with Ad-FOXO1_CA_ localized predominantly in the nucleus and Ad-FOXO1_ΔDB_ having diffuse localization (**Fig. 5B**). Transduction with Ad-FOXO1_CA_ for 48 hours led to a significant reduction in cell size compared to empty vector, while Ad-FOXO1_ΔDB_ had no effect (**Fig. 5C**). This effect on cell size was preceded by induction of autophagy gene expression and increased autophagic flux 24 hours post-transduction (**Fig. 5D-I**). To test whether the effect of FoxO1 activation on cell size was autophagy-dependent, we treated cells with 3-methyladenine (3-MA, inhibitor of autophagosome biogenesis) during viral transduction. 3-MA reduced autophagic flux by half compared to Ad-FOXO1_CA_ alone (**Fig. 5J-K**) and blunted the FoxO1_CA_-dependent reduction of cell size (**Fig. 5L**). These collective data indicate that FoxO1-dependent autophagy mediates the regression of cardiomyocyte hypertrophy upon removal of growth-factor-containing fed python plasma.

**Figure 5.**
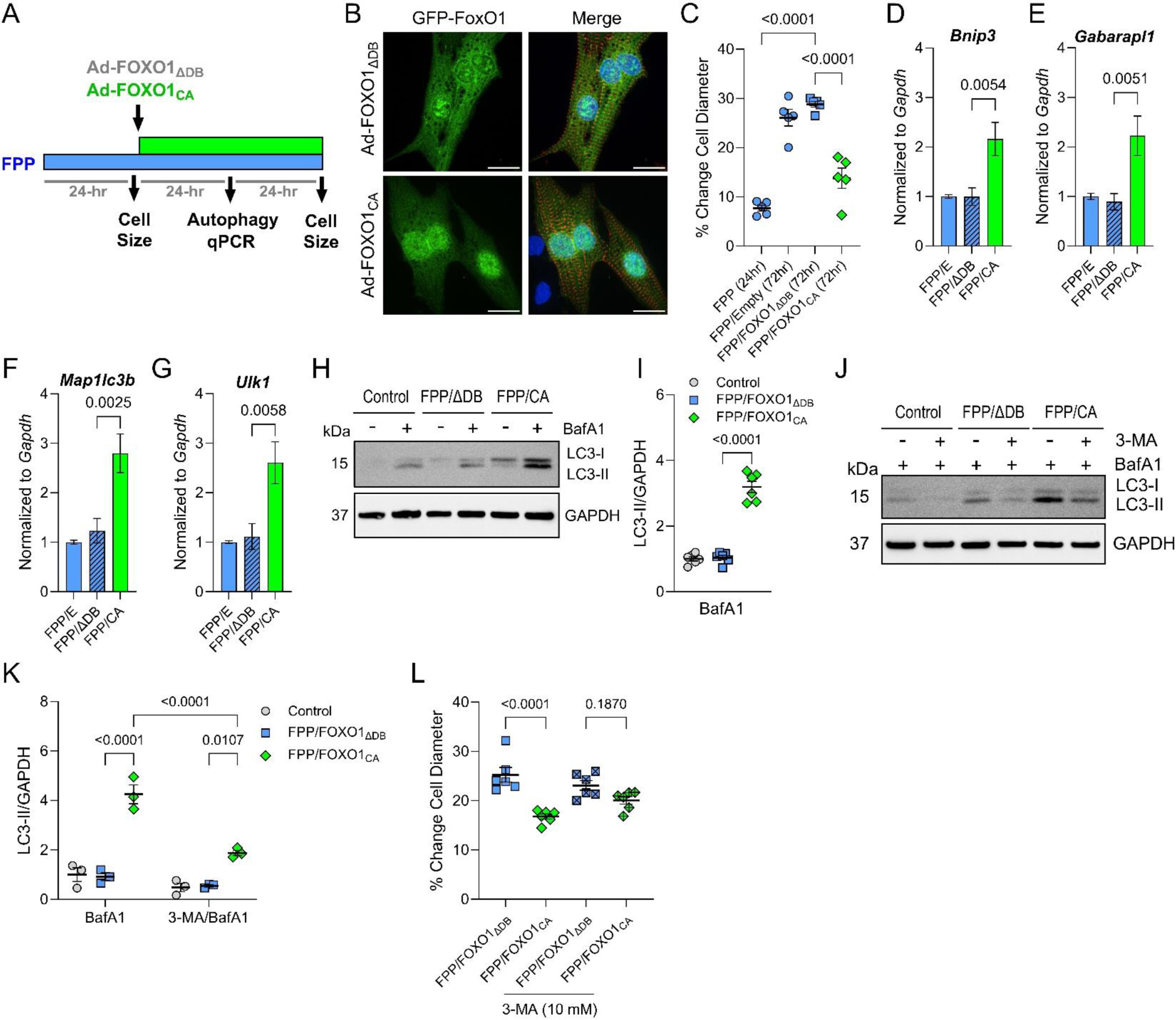
FoxO1 activation stimulates regression of fed python plasma-induced hypertrophy through autophagy. **A.** Experimental paradigm for FoxO1 constitutive activation. **B.** Representative immunofluorescence microscopy images showing localization of adenovirally expressed GFP-FOXO1_ΔDB_ and GFP-FOXO1_CA_ counterstained with α-actinin; 100X magnification, scale bars = 15 µm. **C.** Percent change in cell particle diameter compared to control; n = 10,000-30,000 cells/replicate, 5 biological replicates/group. **D-G.** qPCR analysis for the autophagy genes *Bnip3* (D)*, Gabarapl1* (E)*, Map1lc3b* (F), and *Ulk1* (G) normalized to *Gapdh*; n = 6/group. **H-I.** Representative immunoblot (H) and normalized expression of (I) LC3-II with and without BafA1; n = 6/group. **J.** Representative immunoblot for LC3B and GAPDH in NRVMs exposed to BafA1 ± 10 mM 3-MA. **K.** LC3-II expression normalized to GAPDH. n = 3/group, two-way ANOVA. **L.** Percent change in particle diameter compared to control ± 3-methyl adenine (3-MA); n = 10,000-30,000 cells/replicate, 6 biological replicates/group. For all except K, data were analyzed by one-way ANOVA.

### FoxO1 is activated during regression in mammalian models of reversible cardiac hypertrophy

To determine whether FoxO1 activation was a conserved feature of reversible physiological hypertrophy, we next studied two mouse models of hypertrophy regression from exercise and pregnancy. Chronic exercise was modeled by introduction of a running wheel into the cages for four weeks, which we have previously shown induces hypertrophy^22^. Detraining was initiated by simply removing the wheel. The normalized heart mass in this cohort significantly regressed at 7 days post-running (7DPR). Here, we found that regression correlated with a robust decrease in FoxO1 phosphorylation, indicating increased activity (**Fig. 6A-B**). Moreover, expression of the FoxO1 target genes *Bnip3, Map1lc3b,* and *Ulk1* also increased at 7DPR (**Fig. 6C-E**). Pregnancy in mice also resulted in cardiac hypertrophy that was significantly regressed by one-week post-partum in the absence of lactation (**Fig. 6F**). As in the other models, FoxO1 phosphorylation decreased at one-week post-partum, which correlated with increased expression of *Bnip3* and *Ulk1* (**Fig. 6G-K**).

**Figure 6.**
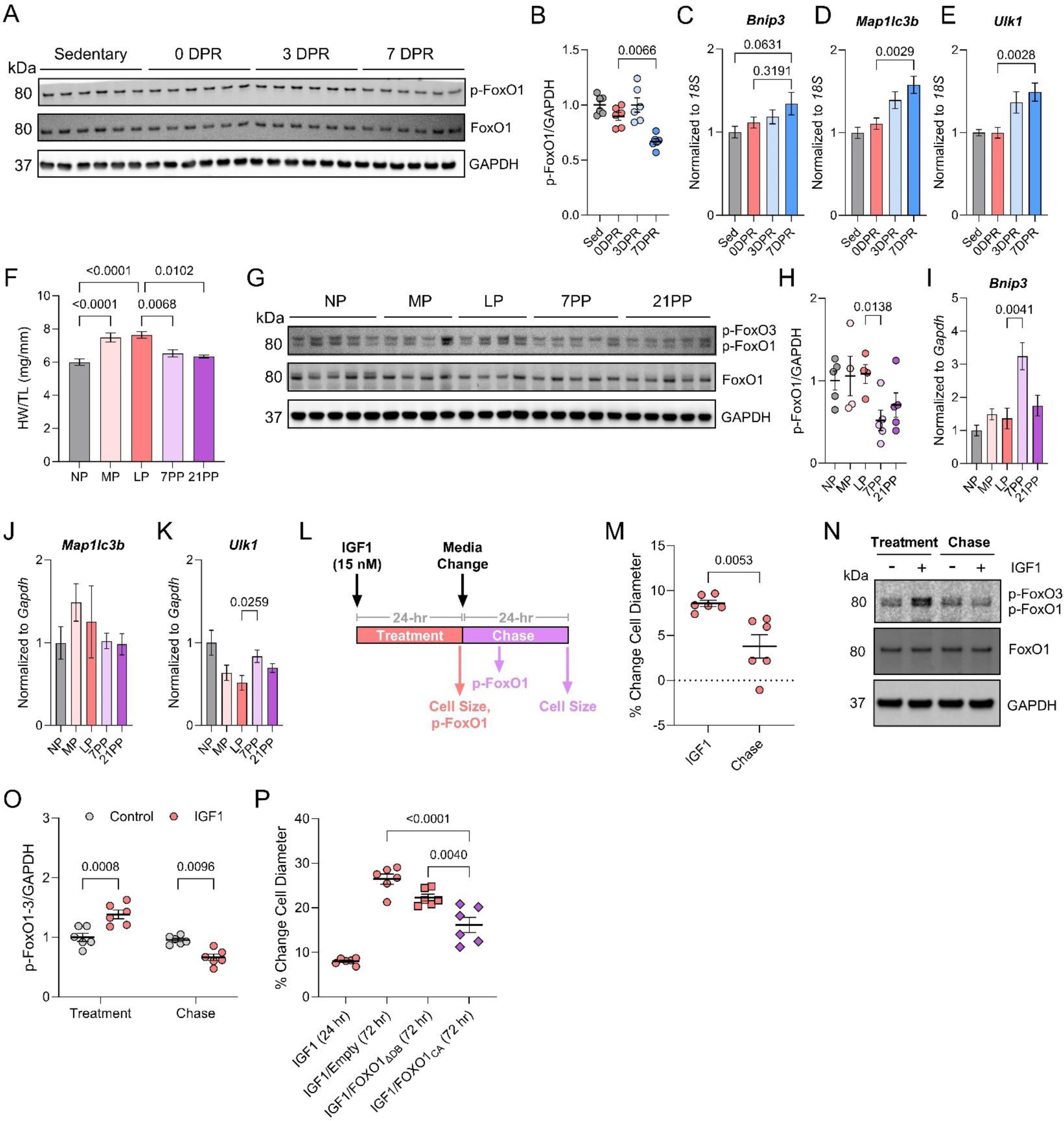
FoxO1 is activated during regression in mammalian models of reversible physiological cardiac hypertrophy. **A.** Western blot for p-FoxO1 (T24), FoxO1, and GAPDH in sedentary and exercised mice; DPR = days post-running. **B.** p-FoxO1 expression normalized to GAPDH in the exercise paradigm. **C-E.** qPCR analysis for *Bnip3* (C)*, Map1lc3b* (D), and *Ulk1* (E) expression normalized to *18S*; for B-E, n = 6/group; one-way ANOVA. **F.** Heart weight normalized to tibia length for non-pregnant (NP), middle pregnant (MP, 13 days), late pregnant (LP, 17 days), 7 days post-partum (7PP), and 21 days post-partum (21PP). Post-partum mice were non-lactating, as pups were immediately removed from the cage following birth. n = 14 NP, 11 MP, 9 LP, 16 7PP, and 7 21PP. **G.** Western blot for p-FoxO1 (T24), FoxO1, and GAPDH in NP, MP, LP, 7PP, and 21PP mice. **H.** p-FoxO1 expression normalized to GAPDH in the pregnancy paradigm. **I-K.** qPCR analysis for *Bnip3* (I)*, Map1lc3b* (J), and *Ulk1* (K) expression normalized to *Gapdh;* for H-K, n = 5 NP, 4 MP, 4 LP, 5 7PP, and 5 21PP; one-way ANOVA. **M.** % change in NRVM cell particle diameter after 24 hours with 15 nM IGF-1 or 24 hours after removal of IGF-1; n = 3/group; two-tailed t-test. **N.** Representative western blot for p-FoxO1, FoxO1, and GAPDH in NRVMs treated with IGF-1 and 6 hours after IGF-1 removal. **O.** p-FoxO1 normalized to total FoxO1; n = 6/group, two-way ANOVA. **P.** % change in NRVM cell particle diameter after IGF-1 with Ad-Empty, Ad-FOXO1_ΔDB_, or Ad-FOXO1_CA_ added at 20 MOI for 48 hours; n = 6/group, one-way ANOVA.

To further extend these findings, we modeled physiological cardiac hypertrophy *in vitro* using NRVMs treated with insulin-like growth factor-1 (IGF1) (**Fig. 6L**). IGF1 is an established agonist for physiological muscle growth and its circulating levels and bioactivity have previously been found to increase with exercise and pregnancy^23–25^. 24 hours of IGF1 exposure caused hypertrophy in NRVMs, which significantly regressed 24 hours after withdrawal (**Fig. 6M**). Hypertrophy and regression in this setting coincided with phosphorylation and de-phosphorylation of both FoxO1 and FoxO3 (**Fig. 6N-O**). Finally, we activated FoxO1 via Ad-FOXO1_CA_ transduction in the presence of IGF1 and found that the IGF1-dependent increase in cell size was partially suppressed (**Fig. 6P**). These collective data support a role for FoxO1 in regression of physiological cardiac hypertrophy in mammals.

## DISCUSSION

Burmese pythons evolved to employ a sit-and-wait predation strategy that is associated with extended periods of caloric restriction (sometimes lasting more than a year) interspersed with consumption of large meals^11^. The extreme fasts combined with the typically large size of their prey precipitates dynamic organ remodeling during digestion, where many organs including the heart hypertrophy to process meal contents, and then rapidly regress during fasting^8–11,26^. The molecular mechanisms underlying the regression of cardiac mass that occurs in pythons following digestion had never previously been explored. Here, we performed a comprehensive analysis of the Burmese python ventricular transcriptome and proteome remodeling throughout digestion, which revealed that FoxO1 activity and autophagy were suppressed during hypertrophy and re-activated with regression. Follow-up *in vitro* examinations using fed python plasma on cultured mammalian cardiomyocytes showed that FoxO1 inhibition prevented regression, while activation of FoxO1 stimulated regression. To examine if this mechanism extended to mammals, we studied mouse models of reversible physiological hypertrophy from exercise and pregnancy. Here also, FoxO1 activity and autophagy gene expression increased during regression, suggesting FoxO1-dependent autophagy may be a conserved mechanism of regression across species.

The involvement of FoxO transcription factors in regulating cardiac and skeletal muscle mass has been described in various other settings. Indeed, FoxO1 and FoxO3 are considered the master transcriptional regulators of skeletal muscle atrophy through induction of proteolytic gene expression^18,27–29^. Earlier work studying FoxOs in NRVMs showed that nutrient starvation induced FoxO1 and FoxO3 activation, leading to increased gene expression of *Atg12*, *Gabarapl1,* and *Map1lc3b*, increased autophagic flux, and FoxO/autophagy-dependent reduction of cell size^30^. FoxOs were also found to be key regulators of mechanical-unloading induced regression of cardiac mass through the atrophic gene *Bnip3*, and this effect was BNIP3/autophagy-dependent^31,32^. Our finding that *BNIP3* is one of the most significantly differentially regulated genes in the python heart during digestion suggests it is also an important factor in this setting. *Bnip3* expression also increased in the post-exercise and post-partum heart suggesting it is a broadly conserved factor upregulated during cardiac regression in physiological settings and not necessarily restricted to a role in atrophy. FoxO activation has also been previously shown to suppress cardiomyocyte hypertrophy in response to hemodynamic stress, neurohumoral factors, and toxic substances in cell and animal models^33–35^. The findings presented herein suggest that reversal of physiological cardiac hypertrophy in pythons and mammals is accomplished through a similar strategy. While it is commonly believed that pathological remodeling is irreversible, we have previously shown that regression of pathological hypertrophy can occur in mice and humans and is associated with improved cardiac function and activation of autophagy^36,37^. However, it is not known whether FoxO1 mediates the regression of hypertrophy in these settings. Future studies investigating whether FoxO1 regulates this response and the therapeutic potential of FoxO1 activation to reverse pathological remodeling in heart disease will be valuable.

Our *in vitro* studies with fed-python plasma and IGF1 exposure in NRVMs support that activation of FoxO1 is a passive response to a reduction in pro-growth signaling induced by circulating factors during early digestion. Since FoxO1 is a direct substrate of Akt^38^, which is activated by hypertrophy-inducing circulating factors to drive muscle growth, the re-activation of FoxO1 with reduced Akt stimulation is a convincing hypothesis for the mechanism underlying regression of physiological hypertrophy. In this vein, Akt, and its upstream regulator PI3K, are positioned as the master controllers of protein homeostasis during physiological cardiac growth, activating protein synthesis and inhibiting FoxO1 to facilitate hypertrophy. When the stimulation of PI3K/Akt wanes after peak digestion in Burmese pythons, the downstream inhibition of FoxO1 is relieved and regression is initiated. Activation of PI3K/Akt signaling has also been described as the primary mechanism driving physiological cardiac hypertrophy in response to exercise and pregnancy in mammals^13,39–42^.

Nevertheless, the data presented herein do not rule out the possibility that an as-yet unidentified atrophic circulating factor could additionally contribute to cardiac mass regression in pythons. The presence of such a factor would help to explain the rate of cardiac mass regression, which seems unlikely to solely be due to a decline in growth signaling. Future studies should focus on identifying circulating lipid and protein factors in 6DPF plasma and exploring their potential to induce regression.

One interesting observation regarding the python cardiac muscle mass regression is that it is not progressive and appears only to occur until a certain baseline is reached without extending into atrophy. In addition, while constitutive activation of FoxOs in the mammalian heart leads to atrophy^43^, FoxO1 and autophagy activity markers are relatively high in the fasted python heart, which maintains a stable mass. Unlike other protein degradation mechanisms, autophagy is an efficient process wherein the amino acids produced through proteolysis are recycled^44,45^. It may be that the python heart has evolved with particularly efficient autophagy regulation that allows it to continually turnover and replace proteins during fasting without significant detriment to muscle mass. Future examinations of python protein turnover mechanisms and how they are impacted during extended fasts will be extremely valuable. Interestingly, results from one previous study found that total muscle mass in Ball pythons (*Python regius*) is hardly affected by a prolonged fast of 168 days^46^. It is likely that this remarkable muscle maintenance during fasting gave pythons an evolutionary advantage for surviving when food was scarce. Understanding how pythons achieve this apparent protection from atrophy at the molecular scale is an important area of future research. Such studies could yield therapeutic targets for preventing cardiac atrophy in humans, as seen in conditions like prolonged bedrest and spaceflight^47^.

In summary, we have identified FoxO1 activation during regression of adaptive physiological cardiac hypertrophy as a conserved molecular feature from pythons to mammals. The potential of FoxO1 as an evolutionarily conserved factor regulating cardiac mass regression is further evidenced by the high sequence conservation in crucial functional and regulatory domains across species^48^. Our collective data support that activation of FoxO1, and subsequent autophagy induction, is a key factor driving cardiac reverse remodeling in physiological settings.

## AUTHOR CONTRIBUTIONS

TGM and LAL designed the study. TGM, DRH, SJL, YT, CCB, CC, and EC performed the experiments. TGM, DRH, and CCB analyzed the data. TGM and DRH designed the figures. TGM and LAL wrote the manuscript with input from all authors.

## ACKNOWLEDGEMENTS

We thank Dr. Massimo Buvoli and Dr. Kristen Bjorkman for their helpful comments on an earlier version of this manuscript. We also thank Aaron Rothchild and the CU Boulder Office of Animal Resources for their assistance with python cohort husbandry. This study was supported by the National Institutes of Health (R01GM029090 to LAL; T32HL007822 and F32HL170637 to TGM.

## METHODS

### Resource Availability

#### Lead Contact

Further information requests should be directed to the corresponding author, Dr. Leslie Leinwand

#### Materials Availability

Unique reagents generated in this study will be made available from the corresponding author upon reasonable request.

#### Data Availability

All study data are included in the article text or associated Supplementary files. The raw RNA sequencing data is deposited in the Gene Expression Omnibus repository (will be uploaded prior to publication). The raw proteomics data is deposited in the PRIDE repository (will be uploaded prior to publication). Reagents and supporting data will be made available from the corresponding author upon reasonable request.

### Ethical Approval

All protocols and procedures involving mice, rats, and pythons were approved by the Institutional Animal Care and Use Committee of the University of Colorado Boulder.

### Animal Models

#### Burmese Python Feeding Model

Captive-bred Burmese pythons (*Python molurus bivittatus*) were purchased from Bob Clark Reptiles, Oklahoma City, OK. The pythons weighed ∼2kg for the RNA sequencing and proteomics studies (n = 6) and ∼250g for full feeding cohort (n = 28). Pythons were fasted for 28 days and then fed a rodent meal consisting of 25% python body weight. Pythons were humanely euthanized under deep isoflurane-induced anesthesia at Fasted (28 day fast), 0.5-, 1-, 2-, 3-, 6-, and 10-days post-fed (DPF) and the ventricle and plasma were collected. The plasma was heat inactivated at 55 °C for 30 minutes for later use in *in vitro* hypertrophy studies. The ventricle tissue was snap frozen in liquid nitrogen and stored at -80 °C.

#### Mouse Exercise Model

Exercise wheels were introduced into the cages of 8–10-week-old C57BL/6J mice. Mice engaged in voluntary cage-wheel running for four weeks. Hearts from mice after four weeks of running were collected as the peak hypertrophy timepoint (0 days post running, DPR). To examine cardiac regression after hypertrophy the exercise wheels were removed from the cages after the month-long exercise period. Hearts from mice in the regression groups were collected at 3 and 7 DPR. Reduction in cardiac mass was significant by 7 DPR. Hearts from sedentary age-matched controls were also collected. Measurements of heart weight normalized to body weight were used to assess hypertrophy and regression.

#### Mouse Pregnancy and postpartum Model

Virgin female C57BL/6J mice, aged 8-10 weeks, were mated with male mice. Successful copulation was confirmed by retrieving the copulatory plug. The mice were then weighed twice weekly, with those showing an increase in body weight at 13 days post-copulation considered pregnant. Mice at this stage were euthanized to collect their hearts to study molecular changes during mid-pregnancy (MP). 17 days post-copulation was designated as late pregnancy (LP). After parturition, the pups were removed from the cage to stop lactation, as lactation is known to further increase pregnancy-induced cardiac hypertrophy^49^. The non-lactating mice were then euthanized at 7 days and 21 days post-partum (PP) for heart collection. Age-matched virgin female mice served as non-pregnant control (NP). Given the expected increase in body weight during pregnancy, heart weight was normalized to tibia length for this study.

### Plasmid Construction and Adenovirus Generation

The plasmids pcDNA-GFP-FKHR, pcDNA-GFP-FKHR-AAA, and pcDNA3-Flag-FKHR-ΔDB-AAA (Addgene 9022, 9023, 10694) were kind gifts from Dr. William Sellers. The EGFP-LC3B plasmid (Addgene 11546) was a kind gift from Dr. Karla Kirkegaard. The open reading frames of the genes of interest were subcloned into pShuttle-CMV using restriction digest cloning, resulting in pShuttle-CMV-EGFP-LC3B, -GFP-FOXO1_WT_, -GFP-FOXO1_CA_ (constitutively active: T24A, S256A, S319A), and -GFP-FOXO1_ΔDB_ (DNA binding domain deficient version of CA: T24A, S256A, S319A, Δ208-220). Plasmids were then transformed into pAdEasy-competent bacteria (BJ5183) and ultimately transfected into HEK293 cells for generation and amplification of adenovirus as previously described^50^.

### Cardiomyocyte Isolation and Culture

Cardiomyocytes were isolated from 1-2 day-old Sprague Dawley rat pups as previously described^51^. Purified ventricular cardiomyocytes were plated at 500,000 cells/well on 6-well plates or 250,000 cells/well on 12-well plates for all experiments. 24 hours after isolation, the wells were washed twice and then left to incubate in experimental media (MEM, 20 mM HEPES, 20 µM Brdu, 2 µg/mL vitamin B12, 50 U/mL PCN-G, 20 U/mL Pen-Strep) for 18 hours prior to the start of the experiment.

#### In vitro Passive Regression Model

PE (10 µM), fasted python plasma (3% final volume), or fed python plasma (FPP, 3% final volume) were added to the media and incubated for 24 hours, at which point collection protein, RNA, or IF staining were performed for the Treatment timepoint. To model Regression, the media was aspirated, wells washed twice with media, and experimental media added (Chase). RNA, protein, and cell size were monitored 24 hours later.

#### FoxO1 Inhibition

The paradigm was the same for this model as the above, except that when media was changed to initiate regression, the FoxO1 inhibitor AS1842856 was added at 2 µM.

#### FoxO1 Activation

Cells were infected with adenoviruses expressing Empty Vector, GFP-FOXO1_WT_, GFP-FOXO1_ΔDB_, or GFP-FOXO1_CA_ at 20 or 40 plaque forming units (PFU) per cell, as detailed in the Figure legends.

RNA and protein were collected after 24 hours, and cell size measured at 48 hours post-infection.

#### Regression of IGF1 Induced Hypertrophy

IGF1 (15 nM) was added in experimental medium and incubated for 24 hours. Cell size was assessed by Coulter Counter after 24 hours to measure hypertrophy. The IGF1-containing medium was then removed, the cells washed twice with medium, and then incubated in experimental medium for 24 hours prior to measuring cell size to monitor regression.

### Cell Size Measurements

To measure cell particle diameter, cells were washed once with PBS and then trypsinized. After lifting, the trypsin was quenched with equal volume addition of 1 mM EDTA, 5% calf serum in PBS. Cells were kept on ice until analyzed. The entire contents of a well were added to 10 mL of Isoton Buffer (Beckman Coulter) and particle diameter measured using a Multisizer 3 Coulter Counter (Beckman Coulter). A 100 µm aperture was used for all experiments. Counting was set to 60 seconds per sample and the mean diameter (representing 10,000-30,000 cells) was used when plotting data.

### RNA Extraction

#### From Ventricular Tissue

Frozen ventricular tissue was homogenized in Trizol and then incubated at room temperature for five minutes. Chloroform was added at 1:5 (v/v), the tubes shaken vigorously, and incubated at room temperature for 15 minutes. Samples were centrifuged at 12,000 RCF for 15 minutes and the upper aqueous layer collected and added to an equal volume of isopropanol. Then samples were incubated at -20 °C for 15 minutes and then centrifuged at 12,000 RCF for 10 minutes to pellet the RNA. The RNA pellet was washed twice with ice-cold 75% ethanol and solubilized in RNase-free water.

#### From Cells

Cells were washed twice with PBS and then collected by scraping in Buffer RLT. RNA was then extracted using the Qiagen RNeasy mini kit according to the manufacturer’s specifications.

### Quantitative RT-PCR

500ng of RNA was used to synthesize complementary DNA (cDNA) with random primers and Superscript III reverse-transcriptase (Invitrogen). 5ng of cDNA was then used for Quantitative RT-PCR with SYBR Green and a Bio-Rad CFX96 Real-Time System. Quantification was performed using the Standard Curve method. The PCR primers used are listed in **Supplemental Table 1**.

### Autophagy Flux Assays

Bafilomycin A1 (BafA1) was used at a final concentration of 50 nM for 6 hours prior to collection of protein lysates or fixing cells for microscopy. To determine autophagic flux in the passive regression model, BafA1 treatment was started after 18 hours of agonist treatment (PE or FPP) to assess flux during hypertrophy development (Treatment). For regression, BafA1 was added 6 hours after the media change (Chase). In the FoxO1 inhibition study, BafA1 was added 18 hours after media change (Chase) with or without the FoxO1-specific inhibitor AS1842856. For experiments using adenoviral expression of *FOXO1*, BafA1 was added 24 hours after infection. Autophagic flux was determined by the level of late-stage autophagosomes using either western blot or immunofluorescence microscopy to assess LC3-II expression. For analysis of microscopy data, the LC3-positive cell area (representing autophagosomes) was quantified using ImageJ. To prevent autophagosome formation, 3-methyl adenine was added to the media at 10 mM final concentration at the time of adenoviral infection and left for 24 or 48 hours before measuring autophagic flux and cell size, respectively.

### *In vitro* Protein Synthesis Assay

After 24 hours of Treatment (fasted plasma, FPP, or PE), Puromycin, or DMSO vehicle control, was added to the media at 1 µM final concentration and the plates returned to the incubator. After 30 minutes, cells were fixed for immunofluorescence microscopy or lysates were collected in RIPA buffer for western blot analysis. Puromycin incorporation was assessed using an anti-puromycin antibody (**Supplemental Table 2**).

### Western Blotting

Cells and ventricular tissue were lysed in RIPA buffer containing Halt protease and phosphatase inhibitors (Thermo) by scraping or pulverizing, respectively. Protein concentration was determined by BCA Assay (Pierce). Protein samples were added to Tris-Glycine SDS Sample Buffer (Novex) containing Bolt Reducing Buffer, boiled for five minutes, and loaded onto Bis-Tris 4-12% gradient gels (Invitrogen). After separation by electrophoresis, proteins were transferred onto nitrocellulose membranes. Blocking was in 5% milk for one hour at room temperature. Primary antibodies (dilutions and vendors in **Supplemental Table 2**) were then added to 5% BSA in TBS-T and incubated overnight at 4 °C. Membranes were washed with TBS-T and then incubated with HRP-linked or near infrared dye (NIR) secondary antibodies (**Supplemental Table 2**) in 5% milk for one hour at room temperature. Membranes were washed with TBS and, for chemiluminescent secondaries, incubated briefly in ECL substrate (Perkin Elmer), and imaged on an ImageQuant LAS 4000 or Azure c600.

### Immunofluorescence Microscopy

This method was adapted from that previously described^52^. Cells were fixed for one minute with ice-cold methanol followed by five minutes in 4% paraformaldehyde. Permeabilization was performed by incubation in 0.5% triton X-100 for 20 minutes and 0.1% triton X-100 for 15 minutes. Coverslips were then incubated in 100 mM glycine for 30 minutes for antigen retrieval, washed three times with PBS, and blocked in 5% BSA for one hour at room temperature. Primary antibodies were added in 5% BSA and incubated overnight at 4 °C. α-actinin (Sigma, A7811, 1:250), LC3B (Cell Signaling, 2755, 1:100), Puromycin (Sigma, Clone 12D10, 1:250). Coverslips were washed with PBS and secondary antibodies (AlexaFluor 488 and/or 568) added at 1:500 in 5% BSA for one hour at room temperature. After a PBS wash, coverslips were mounted with Vectashield (Vector Laboratories), sealed with nail polish, and imaged using either a Nikon Widefield or Nikon Spinning Disc Confocal. Autophagosome area was quantified using ImageJ.

### RNA Sequencing

RNA from the python ventricle (n = 2 per timepoint) was sent to Novogene Corporation (Sacramento, CA) for mRNA enrichment and short-read RNA sequencing at 50 million read-pair depth per sample. Sample quality was assessed using FastQC version 0.11.51^53^ and reads in FASTQ format were cleaned up using Trimmomatic version 0.36^54^ with the following parameters: ILLUMINACLIP:/opt/trimmomatic/0.36/adapters/TruSeq3-PE.fa:2:30:10 \AVGQUAL:24 SLIDINGWINDOW:5:20 MINLEN:75. Trimmed, paired-end reads were mapped to the *Python molurus bivittatus* reference genome using HISAT2 version 2.1.0^55^ with the --sensitive flag. Samtools version 1.8^56^ was used to convert .sam files to sorted and indexed .bam files. The Rsubread version 2.0.1^57^ featureCounts function was used to generate raw counts for each sample. Gene count normalization and differential gene expression analysis was conducted with DESeq2 version 1.36.0^58^. A Likelihood Ratio Test was used to identify genes changing across the post-prandial time course (adjusted p-value < 0.05). Pairwise comparisons between each group were also conducted and the normal shrinkage estimator was used to produce shrunken log fold changes for visualization. Reactome and Gene Ontology (GO) Pathway enrichment of differentially expressed genes was performed using Enrichr and searched against the background gene set of all genes identified in our RNAseq analysis^59^.

### Tissue Preparation for Mass Spectrometry

Python ventricular tissue (n = 2 independently prepped samples per python and 2 pythons per feeding timepoint) was pulverized while frozen and added to solubilization buffer (5% SDS, 0.75% deoxycholate, 50 mM Tris-HCl, pH 8.5). Samples were immediately boiled for 10 minutes and then centrifuged at 12,000 RCF to remove insoluble material. The supernatant was collected, and protein concentration was determined by BCA Assay. Burmese Python heart tissue samples were reduced and alkylated with the addition of 10 mM tris(2-carboxyethylphosphine) (TCEP), 40 mM 2-chloroacetamide, 50 mM Tris-HCl, pH 8.5, boiled at 95°C for 10 minutes and incubated shaking at 2000 rpm at 37°C for 30 minutes. Lysates were digested using the SP3 method^60^. Briefly, 200 µg carboxylate-functionalized speedbeads (Cytiva Life Sciences) were added to approximately 100 µg protein lysate. Addition of acetonitrile to 80% (v/v) induced binding to the beads, then the beads were washed twice with 80% (v/v) ethanol and twice with 100% acetonitrile. Proteins were digested in 50 mM Tris-HCl buffer, pH 8.5, with 1 µg Lys-C/Trypsin (Promega) and incubated at 37°C overnight.

Tryptic peptides were desalted using HLB Oasis 1cc (10mg) cartridges (Waters) according to the manufacturer’s instructions and dried in a speedvac vacuum centrifuge. Approximately 7 µg of tryptic peptide from each tissue sample was labeled with TMT-Pro 16 plex (Thermo Scientific) reagents according to the manufacturer’s instructions. The multiplexed sample was cleaned up with a HLB Oasis 1cc (30mg) cartridge. Approximately 60 µg multiplexed peptides were fractionated with high pH reversed-phase C18 UPLC using a 0.5 mm X 200 mm custom packed UChrom C18 1.8 µm 120Å (nanolcms) column with mobile phases 0.1% (v/v) aqueous ammonia, pH10 in water and acetonitrile (ACN). Peptides were gradient eluted at 20 µL/minute from 2 to 50% ACN in 50 minutes concatenating for a total of 12 fractions using a Waters M-class UPLC (Waters). Peptide fractions were then lyophilized in a speedvac vacuum centrifuge and stored at -20°C until analysis.

### Mass Spectrometry Analyses of TMT Multiplexed Samples

High pH peptide fractions were suspended in 3% (v/v) ACN, 0.1% (v/v) trifluoroacetic acid (TFA) and approximately 1 µg tryptic peptides were directly injected onto a reversed-phase C18 1.7 µm, 130 Å, 75 mm X 250 mm M-class column (Waters), using an Ultimate 3000 nanoUPLC (Thermos Scientific). Peptides were eluted at 300 nL/minute with a gradient from 4% to 25% ACN over 120 minutes then to 40% ACN in 5 minutes and detected using a Q-Exactive HF-X mass spectrometer (Thermo Scientific). Precursor mass spectra (MS1) were acquired at a resolution of 120,000 from 350 to 1500 m/z with an automatic gain control (AGC) target of 3E6 and a maximum injection time of 50 milliseconds. Precursor peptide ion isolation width for MS2 fragment scans was 0.7 m/z with a 0.2 m/z offset, and the top 15 most intense ions were sequenced. All MS2 spectra were acquired at a resolution of 45,000 with higher energy collision dissociation (HCD) at 30% normalized collision energy. An AGC target of 1E5 and 120 milliseconds maximum injection time was used. Dynamic exclusion was set for 20 seconds. Rawfiles were searched against Burmese Python database using MaxQuant v.2.0.3.0. Cysteine carbamidomethylation was considered a fixed modification, while methionine oxidation and protein N-terminal acetylation were searched as variable modifications. All peptide and protein identifications were thresholded at a 1% false discovery rate (FDR). Reporter ion intensities were Cyclic loess normalized and log2 fold changes and p-values were calculated with limma using an R-script. KEGG and Gene Ontology (GO) Pathway enrichment of differentially expressed proteins was performed using Enrichr^59^.

### Statistical Analysis

Statistical analyses and graphical presentation were performed using GraphPad Prism version 9. Comparisons of more than two groups with a single variable were performed using one-way analysis of variance (ANOVA). Comparisons involving two variables were done by two-way ANOVA. When a significant interaction was identified, Tukey’s post-hoc test for multiple pairwise comparisons was employed. Comparisons of two groups were performed by a two-tailed Student’s t-test. The data throughout are presented as the mean ± the standard error of the mean (SEM). A p-value < 0.05 was considered statistically significant.

