## Supplementary Material for "A Conserved Mechanism of Cardiac Hypertrophy Regression through FoxO1"

### SUPPLEMENTAL INFORMATION

**Supplemental Table 1. Primer sequences used for qPCR.**

| Species | Gene | Forward | Reverse |
|---|---|---|---|
| *Python bivittatus* | *BNIP3* | GCTCCTTTGCGCTCTACTATT | AGCCATGCTTCTCTCCATTC |
| *Python bivittatus* | *CTSL* | ACCAATCTCGGTGGCTATTG | ACGCCATGATCCAGTTCTTC |
| *Python bivittatus* | *MAP1LC3B* | CCCGAAGTGAATGCCTCTAATG | GTCTTGCAGTCACCCATGTAAT |
| *Python bivittatus* | *ULK1* | GTGCTGGAAACCTGACTATCTC | AGAGGTCTACGTCCTTCTATGG |
| *Python bivittatus* | *HPRT* | AACAGCGACAAGTCCATTCC | TCTTCATCGTCTTGCCTGTG |
| *Rattus norvegicus* | *Bnip3* | GAGCTGAAATAGACACCCACAG | CCGACTTGACCAATCCCATATC |
| *Rattus norvegicus* | *Map1lc3b* | CCGAAACAGGTCAGGTGTATAG | CCACACTGCTGAGGTGAAA |
| *Rattus norvegicus* | *Ulk1* | GGCTTACAGACTGCCATTGA | GATACCACGCTGGCCTTATAC |
| *Rattus norvegicus* | *Gabarapl1* | TTTGACCTCTGCCCTAATTCC | ATGTCCGTGCGAATGTCTAC |
| *Rattus norvegicus* | *18S* | ACCGCAGCTAGGAATAATGGA | GCCTCAGTTCCGAAAACC |
| *Rattus norvegicus* | *Gapdh* | CATCTCCCTCACAATTCCATCC | GAGGGTGCAGCGAACTTTAT |
| *Mus musculus* | *Bnip3* | CTGAGTAGCAAGTAGAAGCTAAGG | AACTGACCACCCAAGGTAATG |
| *Mus musculus* | *Map1lc3b* | GCGGGTGATTATAGAGCGATAC | CAAGCGCCGTCTGATTATCT |
| *Mus musculus* | *Ulk1* | CAGGGTGGACACATGCTAATAC | CAGCTTGTGGACACTCAGATAC |
| *Mus musculus* | *18S* | GCAATTATTCCCCATGAACG | GGCCTCACTAAACCATCCAA |
| *Mus musculus* | *Gapdh* | TCTCCCTCACAATTTCCATCC | GGGTGCAGCGAACTTTATTG |

**Supplemental Table 2. Antibodies.**

| Antibody | Vendor | Product Number | Dilution |
|---|---|---|---|
| p-FoxO1/3 (T24) | Cell Signaling Technology | 2599 | 1:1000 |
| p-ULK1 (S555) | Cell Signaling Technology | 5869 | 1:1000 |
| LC3B | Cell Signaling Technology | 2775 | 1:1000 |
| p-mTOR | Cell Signaling Technology | 2971 | 1:1000 |
| mTOR | Cell Signaling Technology | 4517 | 1:1000 |
| p-Akt | Cell Signaling Technology | 9271 | 1:1000 |
| Akt | Cell Signaling Technology | 2920 | 1:1000 |
| p-GSK3β | Cell Signaling Technology | 9336 | 1:1000 |
| GSK3β | Cell Signaling Technology | 9832 | 1:1000 |
| GAPDH | Cell Signaling Technology | 2118 | 1:1000 |
| FoxO1 | Cell Signaling Technology | 14952 | 1:1000 |
| Puromycin | Millipore Sigma | 12D10 | 1:1000 |
| HRP Anti-Rb IgG | Cell Signaling Technology | 7074 | 1:2000 |
| HRP Anti-MS IgG | Cell Signaling Technology | 91196 | 1:2000 |
| IR-Dye 800CW | LICOR | 926-32211 | 1:4000 |
| IR-Dye 680RD | LICOR | 926-68070 | 1:4000 |

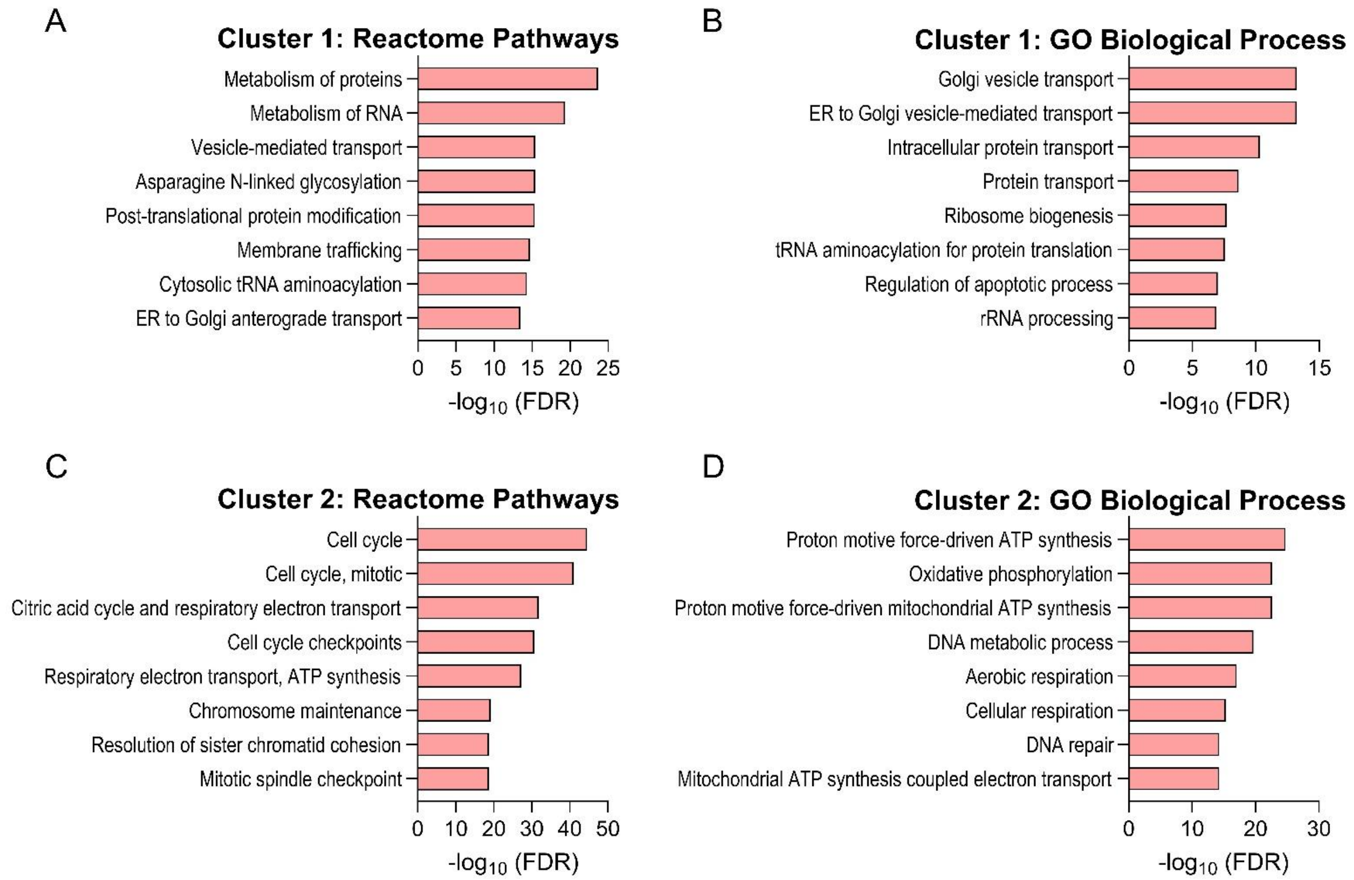


**Supplemental Figure 1. Pathway enrichment for RNAseq Clusters 1 and 2. A-B.** Reactome Pathway (A) and GO Biological Process (B) enrichment for significantly differentially expressed genes in Cluster 1. **C-D.** Reactome Pathway (C) and GO Biological Process (D) enrichment for significantly differentially expressed genes in Cluster 2.

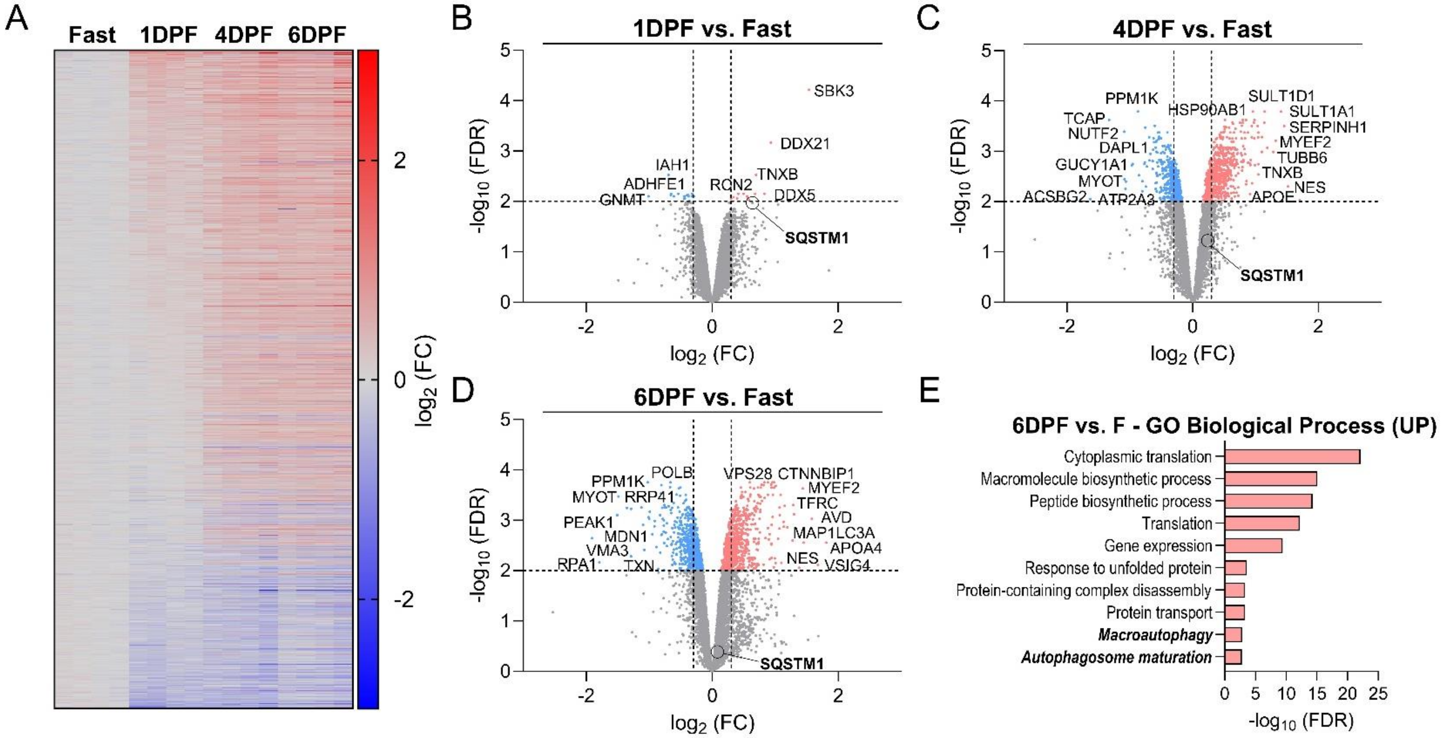


**Supplemental Figure 2. The Burmese python ventricular proteome undergoes dynamic remodeling during digestion. A.** Heat map of significantly differentially expressed ventricular proteins ($p \leq 0.01$, $\log_2$ FC $\geq 0.3$; 1,177 proteins) in 1DPF, 4DPF, and 6DPF pythons compared to Fasted. **B-D.** Volcano plots depicting TMT quantitative proteomics analysis of the python ventricular proteome at 1DPF (B), 4DPF (C), and 6DPF (D) vs. Fasted; y-axis cutoff = 2.0, x-axis cutoffs = -0.3, 0.3; n = 2 pythons per group, 2 separate ventricle samples per snake; SQSTM1 = sequestosome-1 (P62), the increased expression of which corresponds to reduced autophagy activity. **E.** Gene Ontology (GO) Biological Process enrichment of significantly upregulated proteins at 6DPF vs. Fasted.

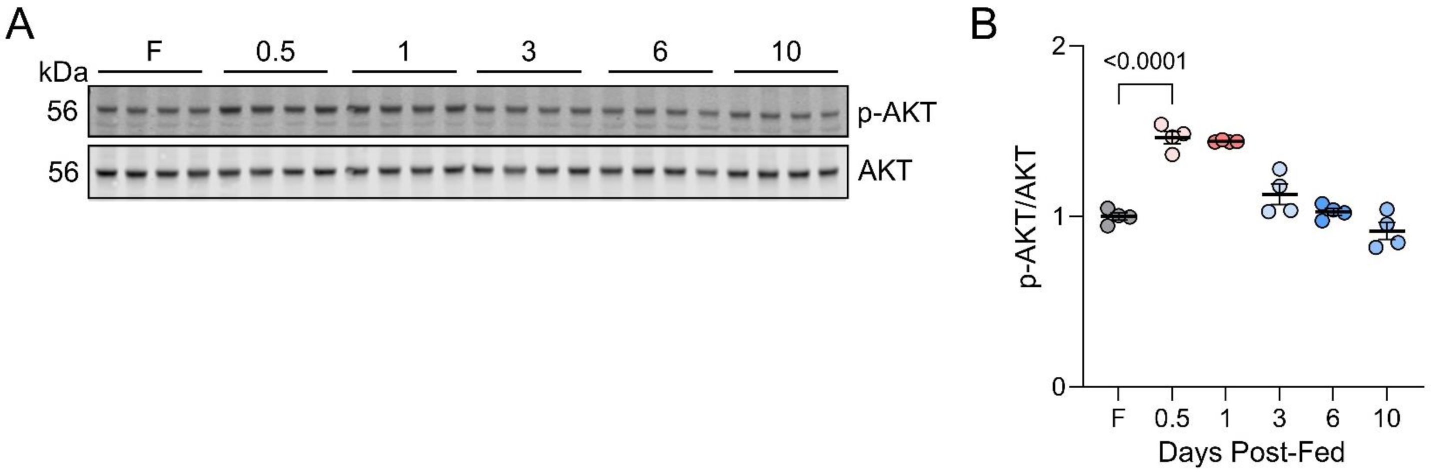


**Supplemental Figure 3. Post-prandial cardiac hypertrophy in Burmese pythons is associated with activation of Akt. A.** Western blot for p-AKT (S473) and total Akt; F = fasted, 0.5, 1, etc. refers to number of days post-feeding. **B.** p-AKT normalized to total AKT; n = 4/group, one-way ANOVA.

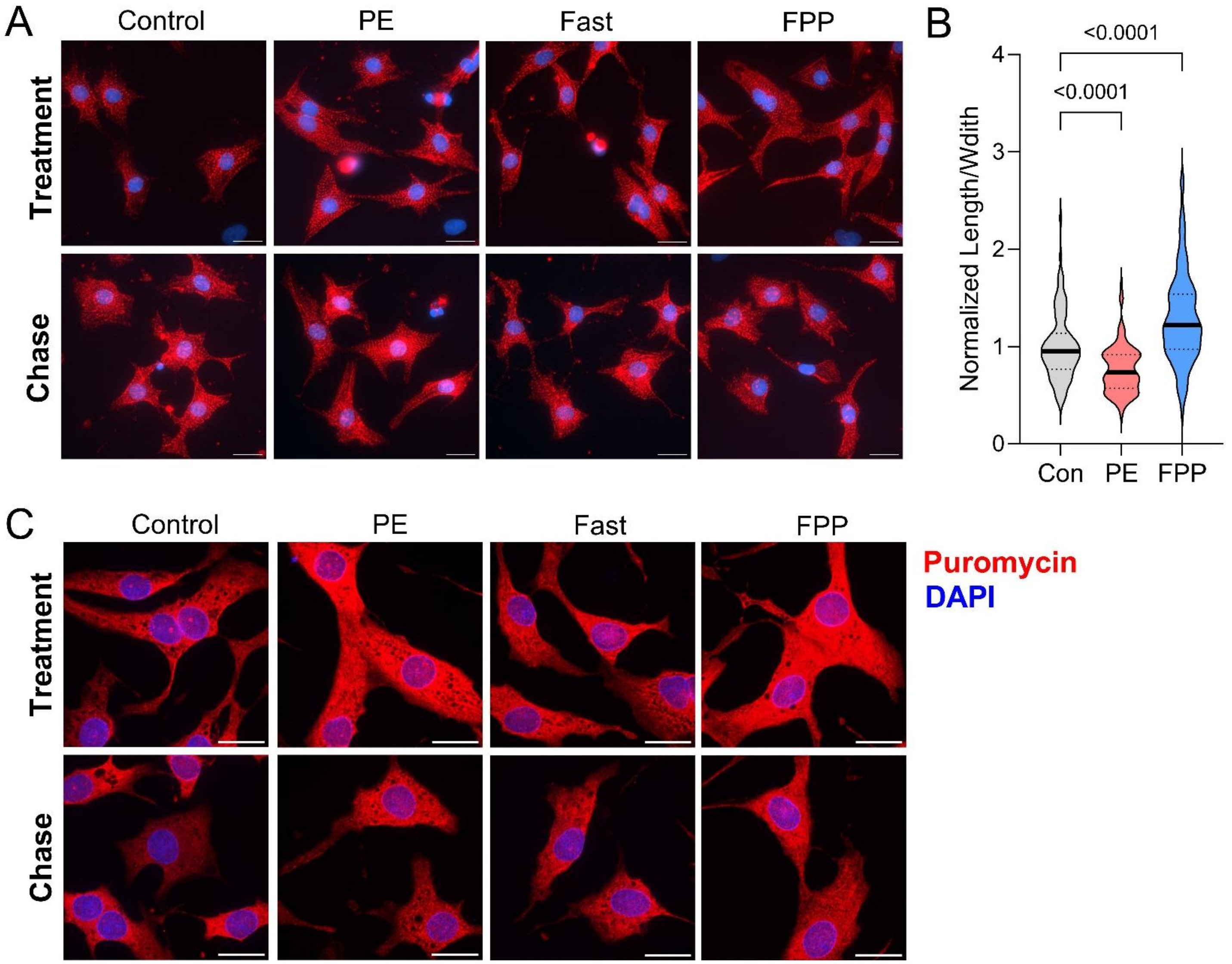


**Supplemental Figure 4. NRVMs treated with fed-python plasma exhibit physiological remodeling and increased protein synthesis. A-B.** Representative immunofluorescence microscopy images for α-actinin (A) and normalized cell length to width measurements (B) with PE or FPP; 40X magnification, scale bars = 15 μm; n = 100 cells per group; one-way ANOVA. **C.** Representative immunofluorescence microscopy images for puromycin; 100X magnification, scale bars = 15 μm.
